# ATP1A3 E815K Disrupts Sodium Homeostasis and Excitability in Human Alternating Hemiplegia of Childhood Neurons

**DOI:** 10.64898/2026.10.08.757730

**Authors:** Mazen Makke, Chungku Lee, Elinor Lazarov, HongJun Rhee, Andreas Ohlenbusch, Lukas Cyganek, Maria-Patapia Zafeiriou, Jutta Gärtner, Jeong Seop Rhee, Hendrik Rosewich

## Abstract

Alternating hemiplegia of childhood (AHC) is a rare neurodevelopmental disorder primarily caused by heterozygous *de novo* mutations in the *ATP1A3* gene, which encodes the neuron-specific catalytic α3 subunit of the P-type Na⁺/K⁺-ATPase. Among these, the E815K variant is the second most frequent ATP1A3 mutation in AHC and is associated with the most severe clinical phenotypes, characterized by frequent epileptic and dystonic episodes and profound developmental impairment, making it a key target for mechanistic study. Although E815K was shown to disrupt pump function in heterologous systems and animal models, its impact on human neurons remain poorly understood. To investigate its disease-relevant pathophysiology, we generated induced pluripotent stem cells (iPSCs) from an AHC patient carrying the E815K mutation and differentiated them into mature neurons. Patient-derived neurons displayed normal differentiation and synaptogenesis, with preserved evoked synaptic transmission. However, α3-specific Na⁺/K⁺ pump currents were nearly abolished, whereas α1 activity remained intact. This selective dysfunction was associated with a depolarized resting membrane potential, reduced input resistance, and diminished intrinsic excitability. Patient neurons generated fewer action potentials in response to depolarizing current steps, displayed altered firing patterns, and showed significantly decreased voltage-gated sodium current densities, while potassium conductance remained unaffected. Taken together, these results indicate that the ATP1A3 E815K mutation disrupts intracellular sodium homeostasis and consequently modifies the excitability of human neurons. Overall, these findings provide mechanistic insight into the molecular pathophysiology underlying this AHC variant and thereby establish a patient-derived neuronal model which serves as a platform for future drug screening approaches and therapeutic investigation.

## Introduction

Alternating Hemiplegia of Childhood (AHC) is a rare, early-onset, and severe neurodevelopmental disorder characterized by recurrent episodes of hemiplegia, dystonia, seizures, developmental delay, cognitive impairment, and a variety of paroxysmal and non-paroxysmal neurological symptoms (Mikati et al. 2000, Heinzen et al. 2012, Rosewich et al. 2012). *De novo* heterozygous mutations in the *ATP1A3* gene were identified as the primary genetic cause of AHC (Heinzen et al. 2012, Rosewich et al. 2012). The *ATP1A3* gene encodes the neuron-specific α3 subunit of the P-type Na⁺/K⁺-ATPase, reported to be the predominant isoform expressed in GABAergic neurons of the brain, and is essential for maintaining electrochemical gradients across the plasma membrane of these cells (Richards et al. 2007, Anderson et al. 2010, Holm et al. 2016, Murata et al. 2020). Among the common *ATP1A3* variants associated with alternating hemiplegia of childhood (AHC), the p.E815K mutation represents the most severe genotype, typically associated with early disease onset, a severe clinical course, frequent epileptic and dystonic episodes, and marked developmental impairment (Rosewich et al. 2012, Panagiotakaki et al. 2015, Viollet et al. 2015, Arystarkhova et al. 2019).

The functional consequences of AHC causing *ATP1A3* mutations, including E815K, have been extensively studied in heterologous expression systems. These studies demonstrated altered Na⁺/K⁺-ATPase pump kinetics, reduced ATPase activity, and impaired ion transport (Arystarkhova et al. 2019, Lazarov et al. 2020, Arystarkhova et al. 2021). Electrophysiological analyses of cortical excitatory neurons differentiated from AHC patient-derived induced pluripotent stem cells (iPSCs) with a G947R mutation confirmed the loss of Na⁺/K⁺-ATPase activity and revealed reduced intrinsic excitability together with increased susceptibility to depolarization (Simmons et al. 2018, Snow et al. 2020) supporting a dominant-negative mechanism underlying the observed neuronal dysfunction.

At the system level, behavioral and electrophysiological investigations in the *Atp1a3* E815K mouse model have revealed spontaneous hemiplegic and dystonic episodes, increased seizure susceptibility, and neurodevelopmental impairments, collectively recapitulating the core clinical manifestations observed in AHC (Helseth et al. 2018, Terrey et al. 2025). Other AHC-associated *Atp1a3* variants, such as D801N and G947R, have also been recapitulated in murine models, which exhibit distinct phenotypic profiles indicative of variant-specific differences in disease severity (Hunanyan et al. 2015, Hawkins et al. 2024, Liu et al. 2024, Terrey et al. 2025). These *in vivo* models have provided critical insights into the electrophysiological and behavioral alterations resulting from impaired function of the α3 subunit. Taken together, these findings underscore the need for integrative, multiscale approaches in patient-relevant models encompassing genetics, biochemistry, cellular modeling, and systems-level neuroscience to elucidate the pathophysiological mechanisms underlying AHC, with particular emphasis on severe genotypes such as E815K.

In this study, we used iPSC-derived GABAergic neurons from an AHC patient carrying the heterozygous *ATP1A3* E815K mutation to investigate the cellular consequences of this variant. Using immunocytochemical and electrophysiological approaches, we demonstrate that the mutant α3 subunit is appropriately expressed in patient-derived neurons and is trafficked into the cell soma and dendrites, and that both synapse formation as well as baseline synaptic transmission are largely preserved. However, the E815K variant causes a profound loss of α3 function, rendering mutant neurons reliant on α1 for Na⁺/K⁺ homeostasis. This results in intracellular ion imbalance, a depolarized resting membrane potential, and, paradoxically, reduced excitability due to a diminished Na⁺ driving force across voltage-gated sodium channels.

## Materials and Methods

### Study subjects and cell lines

The fibroblasts derived induced pluripotent stem cells (iPSCs) used in this study consisted of two clones (UMGi087-A clone 1 and clone 3), generated via lentiviral reprogramming from a 9-year-old male patient diagnosed with AHC. Genetic testing showed that the patient is heterozygous to the mutation 2443G>A (Glu815Lys) in the *ATP1A3* gene. The patient’s first AHC symptoms appeared during the first month of life in the form of dystonic episodes, abnormal eye movements, autonomic dysfunction, and epileptic seizures; hemiplegic episodes appeared at the age of 6 months. With disease progression, the patient exhibited an ataxic gait with muscular hypotonia, developmental delay, dysarthria and intellectual disability.

The control cell lines were generated from two age-matched, unrelated healthy subjects, a male (line UMGi014-C clone 14; generated by Sendai virus), and a female (line UMGi001-A clone 1; generated by lentivirus) (Hanses et al. 2020).

All subjects provided written informed consent for this study, which was approved by the University Medical Center Göttingen.

### Generation of iPSCs

The iPSC lines were generated from fibroblasts isolated from skin biopsies. The skin samples were digested overnight at 37° C in high glucose DMEM containing 20% fetal bovine serum (FBS), 0.25% Collagenase I and 0.05% DNAse-I. Cell suspensions were cultured in T75 flasks and passaged 3 to 4 times before initiating the genetic reprogramming. Fibroblast cells were reprogrammed into iPSCs by overexpression of OCT4, SOX2, KLF4 and c-MYC using the STEMCCA lentivirus.

Between days 29 to 35 post-infection, iPSC colonies appeared and were selected based on their morphology. The colonies were further expanded onto mitomycin C-inactivated mouse embryonic fibroblast (MEF) feeders for several passages using collagenase IV (200 U/ml) in human embryonic stem cell (hESC) medium composted of DMEM/F12 supplemented with 20% Knockout serum replacement, 2mM L-glutamine, 1× nonessential amino acids (NEAA), 50 mM 2-mercaptoethanol, and 10 ng/mL FGF2.

To cultivate the cells in feeder free conditions, cultures were treated with phosphate-buffered saline (PBS)/EDTA solution (0.1% EDTA and 30mM NaCl in PBS) for 3 minutes. The colonies were gently triturated and plated on 3.5 cm dishes coated with either Matrigel or Geltrex Basement Membrane Matrix, in E8 medium supplemented with 10 µM Rock inhibitor (Y-27632). iPSCs were supplemented daily with E8 medium and passaged regularly at 70% confluence, for at least 5 passages before neuronal differentiation was initiated. Reagents are listed in Table S2.

### Reverse transcription PCR for pluripotency markers

Total RNA was reverse transcribed using MuLV reverse transcriptase in a final reaction volume of 20 µl. Each reaction contained 100 ng total RNA in nuclease-free water (2 µl), 8 µl DEPC-treated H2O, 2 µl 10× PCR buffer II, 4 µl 25 mM MgCl₂, 1 µl 100 mM dNTP mix, 1 µl RNase inhibitor (20 U/µl), 1 µl oligo(dT)₁₆ primer (50 µM), and 1 µl MuLV reverse transcriptase (50 U/µl). Reverse transcription was performed in a SensoQuest thermocycler using the following program: 10 min at 22 °C, 50 min at 42 °C, 10 min at 95 °C, followed by a hold at 4 °C. The resulting cDNA was used for PCR amplification of pluripotency-associated transcripts OCT4, SOX2, NANOG, LIN28, FOXD3, and GDF3, as well as the housekeeping gene GAPDH. Each PCR reaction (25 µl) contained 2 µl cDNA template, 14.3 µl DEPC-treated H2O, 5 µl 5× Green GoTaq reaction buffer, 1.6 µl 10 mM dNTP mix, 1 µl forward primer (10 µM), 1 µl reverse primer (10 µM), and 0.1 µl GoTaq DNA polymerase. Gene-specific primer pairs were designed for OCT4, SOX2, NANOG, LIN28, FOXD3, GDF3, and GAPDH and used at identical concentrations (Table S3); cycling parameters were optimized according to primer annealing temperatures while maintaining the same reaction composition (Table S3). PCR was performed on cDNA derived from iPSC lines and control samples, with MEFs serving as negative controls and H₂O as a no-template control to monitor contamination. Amplification products were analyzed by agarose gel electrophoresis to confirm expected amplicon sizes and target expression patterns.

### Flow Cytometry

IPSCs were dissociated by incubation with PBS/EDTA 5 mM for 4 minutes, collected in PBS and singularized by gentle pipetting. Cells were centrifuged for 5 min at 300 g and fixed in 4% PFA for 10 min at RT then washed twice with PBS and blocked in 1% bovine serum albumin (BSA) in PBS for 1-3 days. Cells were then centrifuged at 4° C for 5 min at 500 g, and permeabilized by resuspension in in 1% BSA and 0.1% Triton X in PBS for 10 min on ice. Cells were then centrifuged and the pellet was incubated with the following fluorescently conjugated antibodies diluted in staining solution: anti-OCT3/4-Alexa 647, TRA-1-60-Alexa 488, Hoechst 33342 in 1% BSA with 0.1% Triton-X for 1 hour at 4°C in the dark. After two PBS washes, cells were resuspended in PBS and analyzed using a BD LSRII flow cytometer using FACSDiva software (BD Bioscience).

### Western blots

Cells were harvested in phosphate-buffered saline (PBS), centrifuged for 10 min (210 g, 4°C) and the supernatant was discarded. Cells were resuspended in RIPA-buffer supplemented with Complete protease inhibitor cocktail and incubated on ice for 30 min. Cells were then centrifuged for 15 min at 14000 rpm, 4 °C, and the supernatant was collected. Protein concentration was measured with BCA assay (Interchim BC assay kit) and lysate was mixed with Laemmli buffer and incubated for 10 min at 37 °C. 35 μg of protein was loaded in each lane of polyacrylamide gels. Blotting was performed at room temperature and membranes were blocked with 5% nonfat dried milk in 0.1% Tween 20 in TBS, followed by overnight incubation at 4 °C with primary antibody: anti-OCT4, Pax6, β3 tubulin, ATP1A3, MAP2, SOX2, NANOG, LIN28, FOXD3, or GDF3, with GAPDH as a loading control in blocking solution. After several washes, the membrane was incubated with the HRP conjugated secondary antibodies for 1 hour at RT in blocking solution followed by detection with Lumi-Light or Lumi-Light plus (Roche). Signals were detected by ImageQuant LAS 4000 mini and analyzed with ImageJ.

### Differentiation of iPSCs into GABAergic forebrain neurons

IPSCs were differentiated into GABAergic forebrain neurons as described in Shi et al., (2012) with modifications based on Rhee et al., (2019) to selectively increase the proportion of GABAergic neuronal subtypes. In brief, on day 0, iPSCs were plated at high density onto Geltrex-coated 1/12 well in E8 medium. On day 1, E8 medium was substituted with the neural induction medium containing 3N medium supplemented with 15 µM SB431542 and 500 nM LDN-193189. The 3N medium contained 49% DMEM/F12 Glutamax medium, 49% Neurobasal medium, 1% B27 supplement, 0.5% N2 supplement, 50 µM 2-mercaptoethanol, 100 mM L-glutamine, 0.5% NEAA and 2.5 µg/ml insulin. The neural induction medium was changed every day. Upon neuroepithelial sheet formation (between days 5 and 7), cells were treated with dispase and re-plated in neural induction medium onto Geltrex-coated 3.5cm dishes. Cells were incubated with daily 3N medium changes until rosettes appeared 3 to 4 days after re-plating. The medium was then supplemented with 20 ng/ml FGF2 for 3 days. Between days 12-14, cells were treated with dispase and were further cultivated at 1:2 or 1:3 in 3N medium with medium changes every other day. Between days 17 to 20 neurons accumulated at the rim of the rosettes. Cells were then dissociated with accutase and plated on Geltrex-coated 6-well plates at density of 35,000 cells/cm^2^. Cells were maintained for 4-5 days, with medium change every other day. The cultures were then mitotically inactivated with 5 mM AraC to prevent further proliferation. AraC was removed after 24 h and the cultures were maintained in 3N medium for additional two days. The differentiated neurons were then frozen in cryovials (1 M cells/ml) in freezing medium containing 3N medium, 10% (vol/vol) DMSO, 10 µM Rock inhibitor and 20 ng/ml FGF2. Frozen neurons were thawed and plated onto rat cortical astrocyte micro-islands for further maturation, electrophysiology measurements and immunostaining.

### Autaptic neurons cell culture

To prepare autaptic human neuronal cultures on micro-islands astrocyte layers, we followed a modified version of the protocol previously described by Rhee et al.(Rhee et al. 2019). Astrocytes for the microisland cultures were derived from the cortices of P0 WT mice by digestion using 0.25% trypsin (Gibco) for 15 minutes at 37°C. The cells were then seeded into T75 culture flasks containing DMEM (Gibco) supplemented with 10% fetal bovine serum (FBS, PAA), Mito Serum Extender (Corning), and penicillin/streptomycin (Gibco). The astrocytes were grown for 9-12 days in vitro (DIV). Once the astrocytes reached confluency, they were trypsinized and replated onto 32 mm² glass coverslips, at a density of 30,000 cells per well. The coverslips had been pre-coated with agarose (Sigma) and stamped using a custom-designed stamp to create 400 μm × 400 μm substrate islands. These islands were coated with a solution containing poly-D-lysine (Sigma), acetic acid, and collagen (BD Biosciences). Cryopreserved human neurons (1 ml) were thawed rapidly in a 37°C water bath and diluted in 9 ml of pre-warmed thawing in N3 medium, composed of 49% DMEM/F12, 49% Neurobasal medium, 1% B27 supplement, 1% N2 supplement, 50 μM β-mercaptoethanol, 1 mM L-Glutamax, 1% non-essential amino acids (NEAA), 0.5% FBS, 2 μg/ml doxycycline (Sigma), and 5 μM Y-27632 (Tocris). Cell viability was assessed with Trypan Blue staining. For the autaptic micro-island culture, 15,000 human neurons were seeded per well in a 6-well format onto the pre-cultured astrocyte plates, using either 1.5 or 2.5 mL of N3 medium. The plates were incubated at 37°C for 24 hours. The following day, the entire medium was replaced with 2 mL of freshly conditioned N3 medium. The medium was subsequently changed twice per week until the cells were ready for use. Unless otherwise specified, all materials for cell culture were sourced from Life Technologies.

### Immunofluorescence staining and imaging

To assess the maturation of iPSC-derived GABAergic neurons and the subcellular localization of the α3 subunit, immunocytochemical analyses were performed at days 27–30 of differentiation. Cells were plated onto Poly-L-ornithine/Laminin-coated 12 mm diameter glass coverslips at density of 35,000 cells/cm^2^. Cells were washed twice in PBS (with Ca^2+^ and Mg^2+^) and fixed with 4% formaldehyde in PBS for 8 minutes at room temperature (RT). Cells were washed 3 times with PBS and permeabilized with 0.5% Triton X-100 in PBS for 5 minutes, followed by 5-minute incubation with 0.1% Tween 20 in PBS (PBS-T). Cells were then blocked for 1.5 hours at RT in 3.5% BSA and 0.1% Tween 20 in PBS, and incubated with the primary antibody diluted in blocking solution overnight at 4°C. Secondary antibodies were applied in blocking solution for 45 minutes. The coverslips were washed 3 times with PBS-T and mounted on microscope slides using Prolong Gold antifade with DAPI. Images were taken using an inverted widefield microscope (Axio Imager.M1, Zeiss) using 60x oil-immersion objective and an AxioCam 105 color camera (Zeiss). For the synapse quantification, neurons were fixed in 4% paraformaldehyde in PBS (pH 7.4) for 15 minutes at RT, then washed with PBS. The samples were quenched with 50 mM glycine for 10 minutes and permeabilized using 0.1% Triton X-100 and 10% normal goat serum (NGS) in PBS for 30 minutes at RT. Incubation with the primary antibody was overnight at 4°C in PBS containing 5% NGS and 0.1% Triton X-100. Samples were then washed and incubated with secondary antibodies for 45 minutes at RT followed by extensive PBS washes. Samples were mounted using Aqua-Poly Mount. The primary antibodies for MAP2, SHANK2 and Synapsin 1/2 as well as the corresponding secondary antibodies and their dilutions are in Table S1. Confocal imaging was performed on a Leica SP8 microscope and LAS X software (Leica Microsystems) using a 63x oil objective. Images were analyzed using IMARIS x64 (10.1.1) software. Synapse counting was performed by digital reconstructions to quantify pre-synaptic (Synapsin 1/2) and post-synaptic (SHANK2) puncta located within 1 µm of MAP2-labeled neurons’ soma surfaces. The number of colocalized pre- and post-synaptic signals within 1 µm was also measured. Statistical analysis was performed using GraphPad Prism.

### Electrophysiology

Electrophysiological experiments were conducted using a HEKA EPC-9USB amplifier with PatchMaster software (HEKA Electronics). During voltage whole-cell patch-clamp recordings, cells were held at −70 mV, and EPSCs were evoked by depolarizing the cells from −70 mV to 0 mV. NBQX (HelloBio) and Bicuculline (Tocris) were applied to block AMPA and GABA receptors, respectively. The extracellular solution contained 140 mM NaCl, 2.4 mM KCl, 10 mM HEPES, 10 mM glucose, 4 mM CaCl2, and 4 mM MgCl2 (320 mOsmol/liter), while the patch-pipette solution consisted of 136 mM KCl, 17.8 mM HEPES, 1 mM EGTA, 4.6 mM MgCl2, 4 mM NaATP, 0.3 mM Na2GTP, 15 mM creatine phosphate, and 5 U/ml phosphocreatine kinase, adjusted to pH 7.4 and 315-320 mOsmol/liter. In current-clamp mode, R_input_ was calculated from changes in membrane potential in response to 1-second current injections from −100 to 0 pA in 20 pA increments. Action potentials (APs) were induced by progressively increasing the depolarizing current injections starting from 0 pA in +5 pA steps. All active electrical properties were analyzed from the first trace in which APs were elicited. The external solution for AP recordings was the same as the one used for synaptic current analysis. The patch-pipette solution contained 138 mM K-gluconate, 16.8 mM HEPES, 10 mM NaCl, 1 mM MgCl2·6H2O, 0.25 mM EGTA, 4 mM ATP-Mg^2+^, and 0.3 mM GTP–Na^+^, adjusted to pH 7.4 and ∼320 mOsmol/liter. Voltage-dependent sodium and potassium currents (I_Na_ and I_K_) were evoked by applying 500 ms test pulses from a holding potential of −70 mV to potentials between −80 mV and +70 mV in 10 mV steps. I_K_ was recorded using the same protocol in the presence of 300 nM TTX. I_Na_ was obtained by subtracting I_K_ traces from the total current traces. Total I_K_ was evoked by applying 500 ms depolarizing test pulses from a holding potential of −70 mV to potentials between −80 mV and +70 mV in 10 mV increments, and delayed rectifier I_K_ (I_DR_) was measured under the same protocol in the presence of 1 mM 4AP. A-type I_K_ (I_A_) traces were acquired by subtracting I_DR_ from I_K_. The density of all voltage-gated ion currents was normalized to cell membrane capacitance (pF).

To assess Na^+^/K^+^ pump current density in voltage-clamp mode, outward currents were elicited using 5.4 mM K^+^ and recorded both in the presence and absence of the Na^+^/K^+^ pump inhibitor ouabain (at concentrations of 10, 25, and 100 µM). The internal pipette solution consisted of 30 mM NaCl, 20 mM KCl, 70 mM CsCl, 5 mM MgCl_2_, 5 mM HEPES, 5 mM Na_2_ATP, and 5 mM EGTA. The external bath solution was composed of 60 mM NaCl, 80 mM TEA-Cl, 1 mM MgCl_2_, 2.5 mM CaCl_2_, 5 mM NiCl_2_, 5 mM glucose, and 10 mM HEPES (adjusted to pH 7.4 with CsOH). To evoke pump currents (I_pump_), the bath solution was switched to one containing 60 mM NaCl, 80 mM TEA-Cl, 5.4 mM KCl, 1 mM MgCl_2_, 2.5 mM CaCl_2_, 5 mM NiCl_2_, 5 mM glucose, and 10 mM HEPES (pH 7.4, CsOH). I_pump_ was recorded at a holding potential of −40 mV, and the ouabain-sensitive currents were quantified as the average current over the final 10 seconds of steady-state of each ouabain concentration in the presence of 5.4 mM K^+^. All electrophysiology data were analyzed using Axograph (Ver.1.8.0) software (Axograph Scientific).

## Results

### Generation of Patient-Derived and Control Human iPSCs

AHC is one of the three primary neurological disorders caused by heterozygous missense mutations in the *ATP1A3* gene, which encodes the α3 subunit of the P-type Na⁺/K⁺-ATPase. To investigate the impact of the disease-causing E815K variant on neuronal differentiation and function, fibroblasts were obtained from a 9-year-old male AHC patient and two unrelated healthy individuals (Figure 1A). The study was approved by the institutional review board of the Faculty of Medicine, University of Göttingen, Göttingen, Germany (file reference 15/6/11), and written informed consent was obtained from parents of the participants. Sequencing using the primers specific for *ATP1A3* exon 17-18 (Table S4) confirmed the presence of the heterozygous c.2443G>A (p.E815K) mutation in exon 18 of *ATP1A3* in the patient sample (Figure 1B). This variant is associated with the most severe clinical phenotype of AHC. Control individuals showed no mutations in the *ATP1A3* gene (Figure 1B).

**Figure 1.**
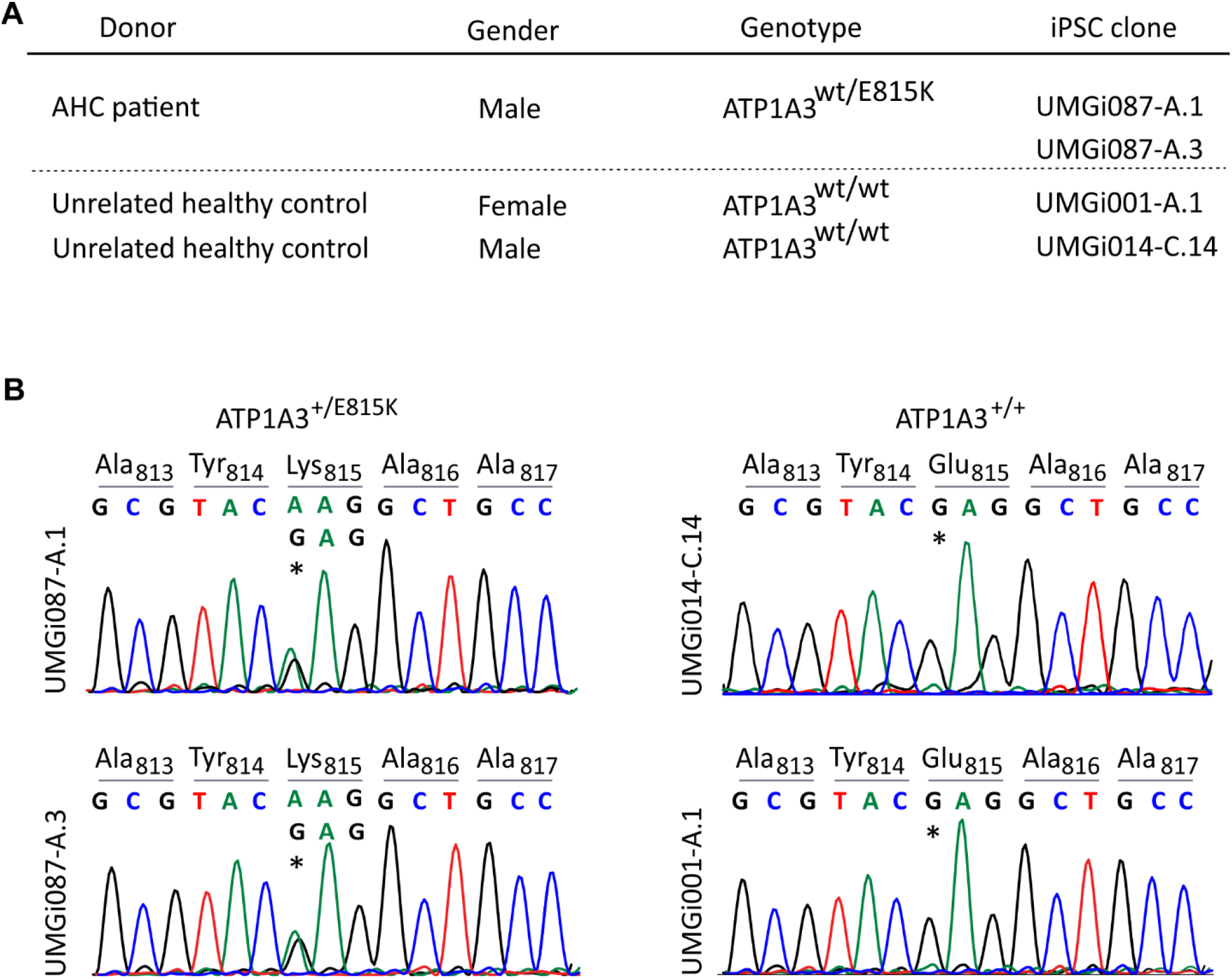
Genetic background of control and AHC patient-generated iPSCs. **(A)** Fibroblasts were collected from one AHC patient and two unrelated healthy individuals (controls). In total, two mutant clones and both control clones were used in the experiments. **(B)** Sanger sequencing results of exon 18 of the *ATP1A3* gene verified the heterozygous missense mutation c.2443G>A in the patient, resulting in the predicted the substitution of Glutamate 815 with Lysine (p.E815K) in approximately half the expressed α3 subunits of the Na^+^/K^+^ pumps. The control lines displayed the wildtype sequence, indicating an intact *ATP1A3* gene exon 18 sequence.

Fibroblasts were reprogrammed into iPSCs, generating two independent clones per genotype: from the AHC patient, UMGi087-A.1 and UMGi087-A.3 were derived via lentiviral reprogramming, while iPSC lines from age-matched healthy controls included UMGi001-A.1 (female; lentivirus) and UMGi014-C.14 (male; Sendai virus). All iPSC lines exhibited normal karyotypic integrity (Figure S1A) and robustly expressed canonical pluripotency markers (Figure S1B, C), indicating successful reprogramming.

### AHC-associated ATP1A3-E815K iPSCs Exhibit Normal Neuronal Differentiation

We differentiated human (iPSCs into neurons over a three-week period (see Materials and Methods) (Figure 2A). Throughout the differentiation process, we monitored the cells for neurite outgrowth (Figure 2B) and performed Western blot analysis on total protein extracts collected at the different stages of differentiation (Figure 2C). In undifferentiated iPSCs (Day 1), we observed a strong expression of the pluripotency marker Oct4, minimal expression of the neuronal maturation marker β3 tubulin, and no detectable signal for the neuronal precursor marker Pax6 (Figure 2C). By the intermediate stage of differentiation (Days 10–13), Oct4 expression was no longer detectable, while Pax6 expression peaked, accompanied by a significant increase in β3 tubulin levels (Figure 2C). At the terminal stage of differentiation (Days 28–31), PAX6 expression declined, whereas β3-tubulin levels further increased, consistent with the predominance of mature neurons in the culture (Figure 2C). Next, we assessed the expression of the Na⁺/K⁺-ATPase α3 subunit in differentiated neurons by Western blot analysis. A distinct immunoreactive band corresponding to the α3 subunit (∼110 kDa) was detected in total protein extracts from differentiated neurons, consistent with its expression pattern in mouse brain tissue. In contrast, no α3 subunit signal was observed in undifferentiated iPSCs or in mouse liver, which served as a negative control, confirming the antibody’s specificity for the α3 subunit (Figure 2D). Immunocytochemistry analysis further revealed that the differentiated neurons expressing β3 tubulin also expressed GABA, a marker for inhibitory neurons, confirming their GABAergic identity (Figure 2E). Additionally, we show that the Na^+^/K^+^ α3 subunit colocalizes to the somato-dendritic marker MAP2 in both patient-derived (E815K mutation) and wild-type (WT) neurons. These findings suggest that the E815K mutation neither impacts neuronal differentiation nor alters the subcellular localization of the α3 subunit in GABAergic neurons. However, whether this mutation affects the functional properties of the Na^+^/K^+^ pump and neuronal activity remains to be determined.

**Figure 2.**
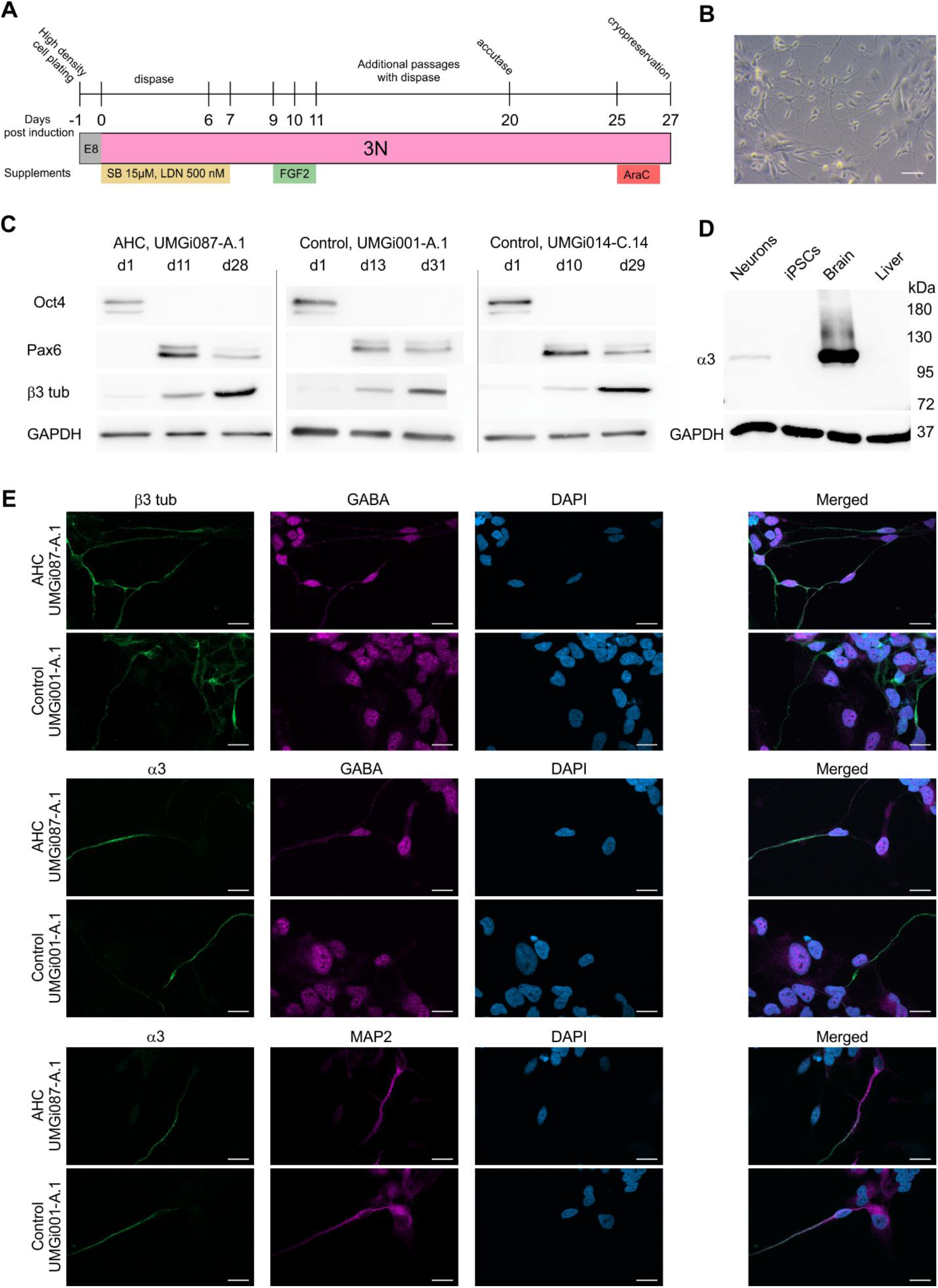
Differentiation of healthy WT and AHC patient derived iPSCs into mature inhibitory neurons. **(A)** Schematic depicting the timeline and handling stages for the differentiation of iPSCs into mature neurons. **(B)** Phase contrast image of differentiated AHC neurons (cell line UMGi087-A.1) prior to cryopreservation, scale bar 50 µm. **(C)** Western blot analysis of iPSCs harvested at different stages of the differentiation. At day 1, the pluripotency marker Oct4 (∼45 kDa) is detected, with only trace amounts of β3-tubulin (∼55 kDa). The expression of the neuronal precursor marker Pax6 (∼43 kDa) is highest at days 10 to 13, coinciding with an increase β3-tubulin expression. At days 28 to 31, Pax6 levels decrease and the neuronal marker β3-tubulin shows its highest expression. **(D)** The differentiated human neurons as well as mouse brain (positive control), but not the undifferentiated iPSCs or mouse liver (negative control), show a band for ATP1A3 at ∼110 kDa (days 24-25, UMGi087-A.1 line). **(E)** Immunostaining of AHC and control neurons at post-induction days 28-31. Shortly after emergence, differentiated neurons express β3-tubulin and GABA. Scale bar 20 µm. **(F)** Inhibitory neurons differentiated from both experimental groups expressed ATP1A3. Scale bar 20 µm. **(G)** Co-staining of ATP1A3 and MAP2 indicates the successful induction of mature inhibitory neurons. ATP1A3 colocalizes to MAP2 positive structures in both healthy WT and AHC patient differentiated neurons. Scale bar 20 µm.

### Synaptogenesis and Synaptic Transmission Are Preserved in AHC Neurons

To evaluate whether the E815K mutation affects synaptogenesis, differentiated neurons were immunostained for the presynaptic marker Synapsin1/2 and the postsynaptic marker SHANK2 (Figure 3A). Quantification of Synapsin1/2-positive puncta showed no significant differences (P = 0.8181) between control (675.5 ± 146.8, n = 6) and patient-derived neurons (734.5 ± 202.2, n = 6) (Figure 3B). Similarly, SHANK2-positive puncta counts were comparable between controls (379.2 ± 72.2) and patient-derived neurons (360.7 ± 89.0) (P = 0.8750) (Figure 3C). Consistent with these individual marker counts, the number of colocalized puncta did not differ significantly (control: 341.8 ± 83.9, n = 6; patient: 332.3 ± 75.0, n = 6; P = 0.9344) (Figure 3D), nor did the relative proportion of colocalized puncta (control: 0.497 ± 0.079, n = 6; patient: 0.502 ± 0.044, n = 6; P = 0.9600) (Figure 3E). These findings indicate that the E815K mutation does not impair synapse formation in patient-derived neurons.

**Figure 3.**
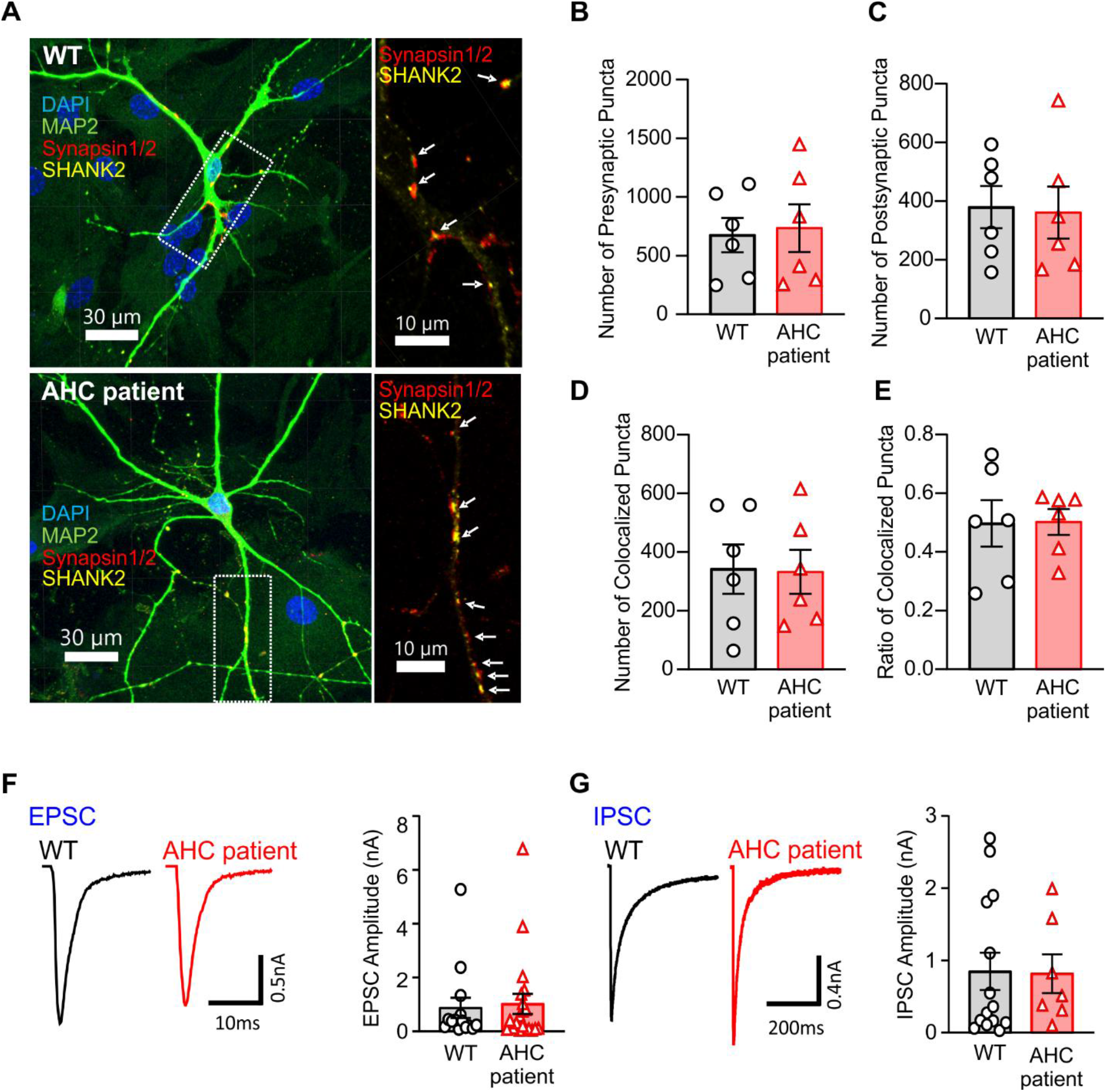
E815K α3 mutant neurons show normal synaptogenesis and evoked synaptic transmissio. **(A)** Confocal images of healthy (WT) and E815K AHC mutant neurons cultured for 8 weeks on mouse astrocyte micro-islands. Cells were stained for the presynaptic marker Synapsin 1/2, the postsynaptic marker SHANK2, the neuronal somato-dendritic marker MAP2, and the nuclear marker DAPI. Rectangles indicate the position of the magnified regions, and arrows point to representative synapses. **(B)** Quantification of presynaptic puncta showing similar counts in WT (n = 6) and AHC patient neurons (n = 6). **(C)** Quantification of postsynaptic puncta showing similar counts in WT (n = 6) and AHC patient neurons (n = 6). **(D)** Quantification of colocalized pre and post synaptic puncta in WT (n = 6) and AHC patient neurons (n = 6). **(E)** Ratio of colocalized pre- and postsynaptic puncta in WT (n = 6) and AHC patient neurons (n = 6). **(F)** Exemplary excitatory postsynaptic currents (EPSCs), and quantification of EPSC amplitude in WT and AHC patient neurons (WT: n = 14; AHC patient: n = 20). **(G)** Exemplary inhibitory postsynaptic currents (IPSC) and quantification of IPSC amplitude in WT and AHC patient neurons (WT: n = 14; AHC patient: n = 7).

Electrophysiological recordings from autaptic neurons showed no significant differences in evoked excitatory postsynaptic current (EPSC) amplitudes between control (0.87 ± 0.38 nA, n = 14) and patient-derived neurons (1.02 ± 0.37 nA, n = 20) (P = 0.7821) (Figure 3F). Similarly, inhibitory postsynaptic current (IPSC) amplitudes were comparable (control: 0.85 ± 0.26 nA, n = 14; patient: 0.82 ± 0.27 nA, n = 7; P = 0.7004) (Figure 3G). These data indicate that both synapse formation and evoked synaptic transmission are intact in neurons carrying the *ATP1A3* E815K mutation.

### ATP1A3 E815K Profoundly Diminishes the α3-Dependent Na⁺/K⁺-ATPase Currents in AHC Patient Neurons

The ATP1A3 E815K mutation has been shown to cause a loss of function in heterologous expression systems, as demonstrated in COS-7 cells (Heinzen et al. 2012). This impairment is further supported by *in vivo* studies in *Atp1a3*^E815K/+^ mutant mice reporting a complete loss of α3 subunit-dependent Na⁺/K⁺-ATPase activity (Helseth et al. 2018). To determine whether the *ATP1A3* E815K mutation similarly impairs the function of the α3 Na⁺/K⁺-ATPase isoform in AHC patient-derived neurons, we quantified residual pump-mediated currents in response to increasing concentrations of ouabain, a specific Na^+^/K^+^ pump blocker, thereby isolating the α3 subunit–dependent component of Na⁺/K⁺-ATPase activity.

Total Na⁺/K⁺ pump currents were measured at a holding potential of −40 mV by increasing extracellular K⁺ from 0 to 5.4 mM, which evoked a step-like inward current that returned to baseline upon K⁺ removal (Figure 4A). Total pump current density was significantly reduced in AHC patient-derived neurons compared with controls (0.262 ± 0.038 pA/pF, n = 9 vs. 0.409 ± 0.042 pA/pF, n = 11; P = 0.0204) (Figure 4B). Subunit-specific contributions were then resolved using differential ouabain sensitivity. Selective inhibition of α3 with 10 µM ouabain revealed a marked reduction in α3-mediated current density in AHC neurons (0.016 ± 0.006 pA/pF, n = 9) relative to controls (0.092 ± 0.021 pA/pF, n = 11), corresponding to a fourfold decrease in its contribution to total pump activity (5.97 ± 1.71% vs. 20.20 ± 3.73%; P = 0.0048) (Figure 4C). In contrast, α1-mediated currents, measured with 100 µM ouabain, were unchanged between groups (0.205 ± 0.033 pA/pF, n = 9 vs. 0.231 ± 0.027 pA/pF, n = 11; P = 0.5464) (Figure 4D), resulting in a larger relative α1 contribution in AHC neurons (75.76 ± 5.11% vs. 54.04 ± 5.75%). The relative ouabain-insensitive currents were similar between groups (Figure 4E). These findings indicate that α3 pump activity is nearly abolished, whereas α1 function remains intact and accounts for most residual Na⁺/K⁺ pump activity in AHC patient-derived neurons. This suggests that the α1 subunit retains normal expression and membrane localization despite the presence of the E815K mutation.

**Figure 4.**
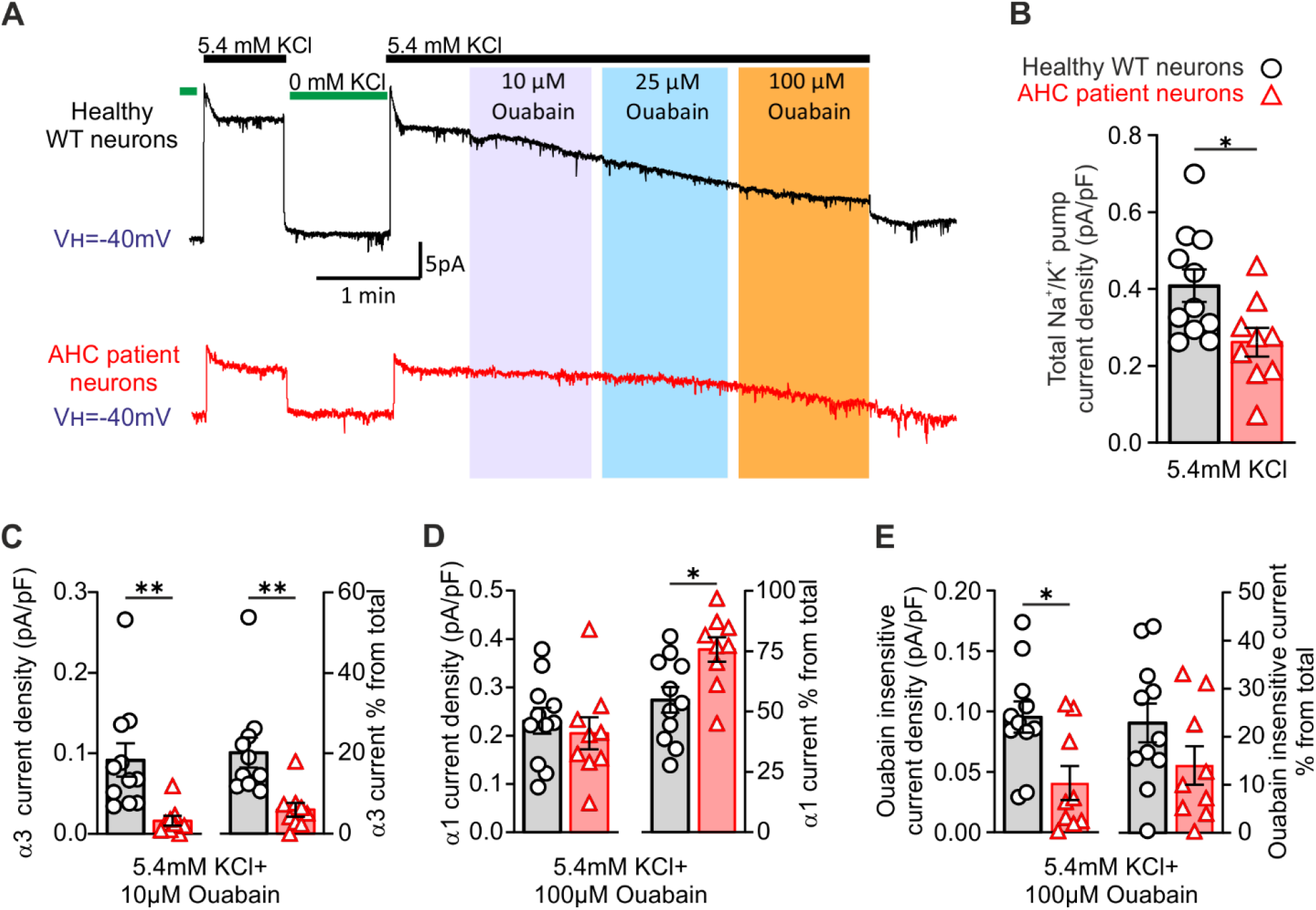
E815K mutation severely reduces α3 subunit activity in patient neurons without affecting α1 subunit function. **(A)** Representative Na^+^/K^+^ pump traces in WT and AHC patient neurons in response to varying concentrations of ouabain. Baseline currents were determined at 0 mM extracellular K^+^ (green line) and pump currents were activated with 5.4 mM extracellular K^+^ (black). **(B)** Quantification of total Na^+^/K^+^ pump currents shows a significant reduction in the AHC patient derived neurons triggered by 5.4mM extracellular KCl (WT: n = 11; AHC patient: n = 9). **(C)** Quantification of the α3 current density and its proportion relative to the total Na^+^/K^+^ current measured in presence of 10 μM ouabain (WT: n = 11; AHC patient: n = 9). **(D)** Quantification of the α1 current density and its proportion relative to the total Na^+^/K^+^ current measured in the presence of 100 μM ouabain (WT: n = 11; AHC patient: n = 9). **(E)** Quantification of the ouabain insensitive current density and its proportion relative to the total Na^+^/K^+^ current measured in the presence of 100 μM ouabain (WT: n = 11; AHC patient: n=9). Data are presented as mean ± SEM. *p<0.05, **p<0.01.

### AHC Patient iPSC-Derived Neurons Exhibit Reduced Excitability Due to *ATP1A3* E815K

Given the critical role of the Na^+^/K^+^ pump in maintaining neuronal ion homeostasis and a stable resting membrane potential, we performed electrophysiological measurements to investigate membrane properties in iPSC-derived neurons from AHC patients with impaired α3 function.

First, the passive membrane properties of iPSC-derived neurons from AHC patients were analyzed. The resting membrane potential (RMP) in the patient group was significantly depolarized at −38.31 ± 1.17 mV (n = 96) compared to −46.16 ± 0.96 mV (n = 91) in the healthy group (P = <0.0001) (Figure 5A), consistent with disrupted ionic homeostasis due to reduced α3 pump activity. Membrane capacitance was similar between groups (24.23 ± 1.77 pF, n = 99 vs. 24.66 ± 1.40 pF, n = 95; P = 0.8507) (Figure 5B), In contrast, the input resistance (Ri_nput_) was significantly lower in patient neurons (1171 ± 47.52 MΩ, n = 90) compared to the healthy group (1305 ± 38.18 MΩ, n = 93) (P = 0.0286) (Figure 5C). This decrease in input resistance indicates increased membrane conductance independent of cell size, likely reflecting altered ion channel activity, and contributing to the reduced neuronal excitability observed in the patient group.

**Figure 5.**
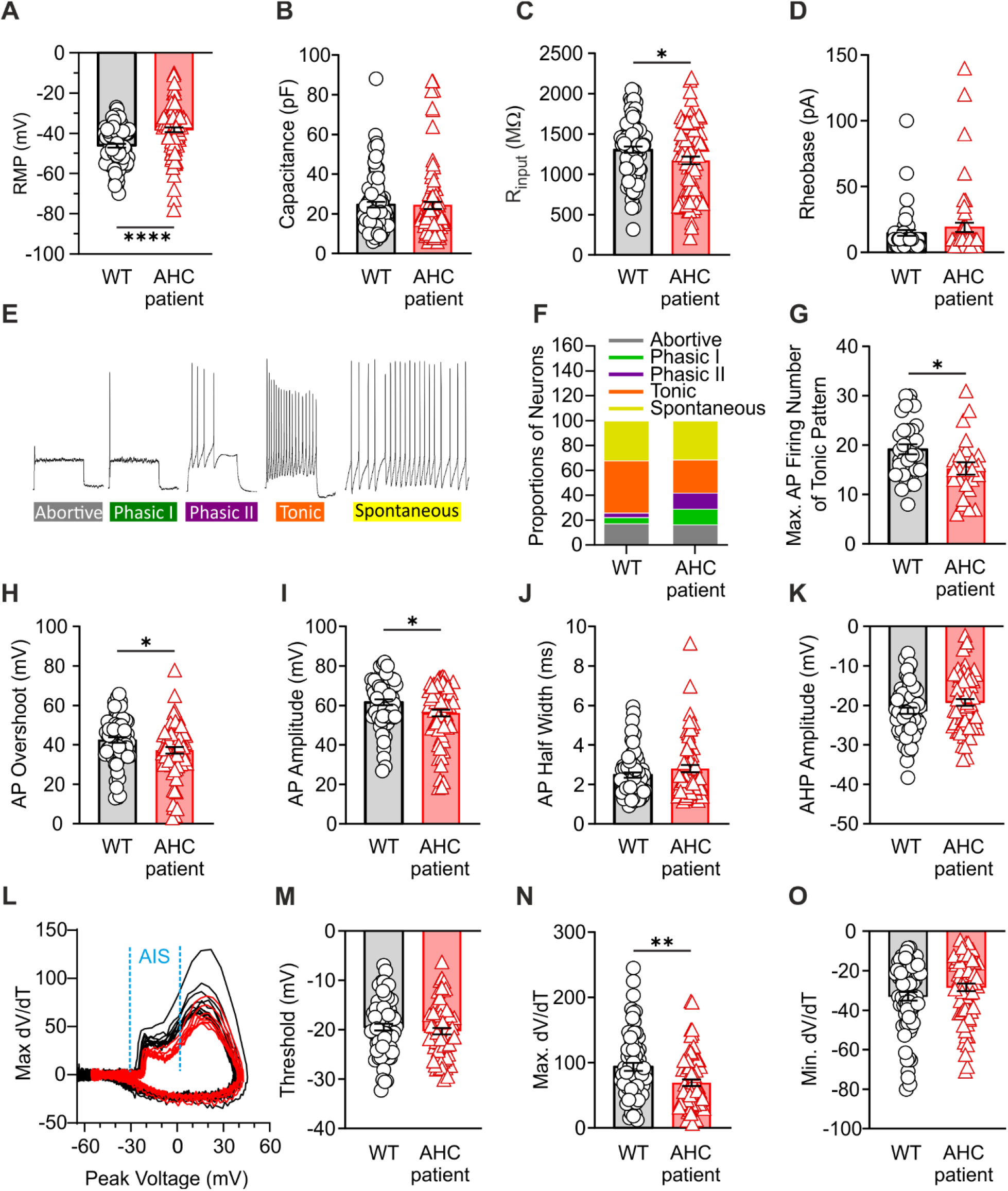
AHC-patient derived neurons are less excitable than WT neurons. **(A)** Average resting membrane potential (RMP) (WT: n = 91; AHC patient: n=96). **(B)** Average membrane capacitance (WT: n = 95; AHC patient: n=99). **(C)** Average input resistance (R_input_) (WT: n = 93; AHC patient: n=90). **(D)** Average rheobase current (WT: n = 54; AHC patient: n=57). **(E)** Representative traces of action potential firing patterns. **(F)** Distribution of action potential firing patterns. **(G)** Average maximum number of action potentials in tonic firing patterns (WT: n = 33; AHC patient: n = 25). **(H)** Average action potential overshoot (WT: n = 68; AHC patient: n = 69). **(I)** Average action potential amplitude (WT: n = 68; AHC patient: n = 69). **(J)** Average action potential half-width (WT: n = 68; AHC patient: n = 69). **(K)** Average after-hyperpolarization (AHP) amplitude (WT: n = 68; AHC patient: n = 69). **(L)** Representative phase plane plots of action potentials, highlighting the axon initial segment (AIS) dominated phase in blue. **(M)** Average threshold voltage for action potential initiation (WT: n = 68; AHC patient: n = 69). **(N)** Average maximum dV/dT of action potential upstroke (WT: n = 68; AHC patient: n = 69). **(O)** Average minimum dV/dT of action potential repolarization (WT: n = 68; AHC patient: n = 69). Data are presented as mean ± SEM. *p<0.05, **p<0.01, ***p<0.001.

Active membrane properties were then assessed. The rheobase did not differ between groups (P= 0.3006) (Figure 5D). Analysis of firing patterns revealed a reduction in tonic firing among patient neurons (26.7% vs. 42% in controls), with corresponding increases in phasic I and phasic II firing patterns (Figure 5E, F). The maximum number of action potentials during tonic firing was significantly lower in patient neurons (15.28 ± 1.23, n = 25) compared to controls (19.18 ± 1.03, n = 33) (P = 0.0177) (Figure 5G). AP overshoot and amplitude were modestly but significantly reduced in patient neurons (overshoot: 37.20 ± 1.68 vs. 42.47 ± 1.44 mV; P = 0.0184; amplitude: 56.11 ± 1.72 vs. 61.60 ± 1.48 mV; P = 0.0170) (Figure 5H, I), while AP half-width (P = 0.1402) and afterhyperpolarization (AHP) amplitude (P = 0.0658) were unchanged (Figure 5J, K). These results suggest that reduced excitability in patient-derived neurons, characterized by depolarized RMP and decreased AP overshoot and amplitude, underlies the shift from tonic to phasic firing, preferentially impairing sustained activity.

Phase-plane analysis of AP kinetics revealed no difference in the AP threshold (P = 0.4108) (Figure 5M), whereas the maximal rate of depolarization (maximum dV/dt) was significantly reduced in patient neurons (P = 0.0026) (Figure 5N), with repolarization kinetics (minimum dV/dt) unchanged (P = 0.1180) (Figure 5O). These findings indicate a slowed AP rising phase, likely reflecting impaired voltage-gated sodium channel function, either due to decreased channel availability or altered local ionic homeostasis at the axon initial segment.

These results suggest that the slowed AP rising phase in AHC patient-derived neurons likely reflects impaired voltage-gated sodium channel function, arising either from decreased channel availability or altered local ionic homeostasis at the axon initial segment, the site of AP initiation (Kole et al. 2008, Jenkins and Bender 2025). The absence of differences in AP threshold and initiation, together with the reduced proportion of tonic firing, fewer APs, and slowed depolarization kinetics, supports the conclusion that altered neuronal excitability in these neurons results from dysfunctional sodium channel gating or reduced sodium current density secondary to disrupted ion homeostasis.

### AHC Patient-Derived Neurons Exhibit Impaired Voltage-Gated Sodium Current Density

To investigate whether the observed decrease in AP amplitude and firing rate in the iPSC-derived neurons from AHC patients is due to disrupted ion homeostasis or altered sodium channel activity, we measured voltage-dependent sodium and potassium currents and calculated their corresponding current densities and voltage-dependent properties.

Sodium current peaks were elicited using 500 ms square pulses applied in 10 mV increments from-80 mV to +70 mV (Figure 6A). The current-voltage (I-V) relationship of sodium current densities exhibited maximal activation between –10 and 0 mV, with no difference in the reversal potential between groups, indicating that the intrinsic biophysical properties of Na⁺ channels remained unaltered (Figure 6B). However, the peak sodium current density was significantly reduced in the AHC patient group (187.3 ± 13.02 pA/pF, n = 46) compared to the healthy group (249.3 ± 16.49 pA/pF, n = 47) (P = 0.0041) (Figure 6A and C), consistent with a functional deficit in sodium channel-mediated excitability.

**Figure 6.**
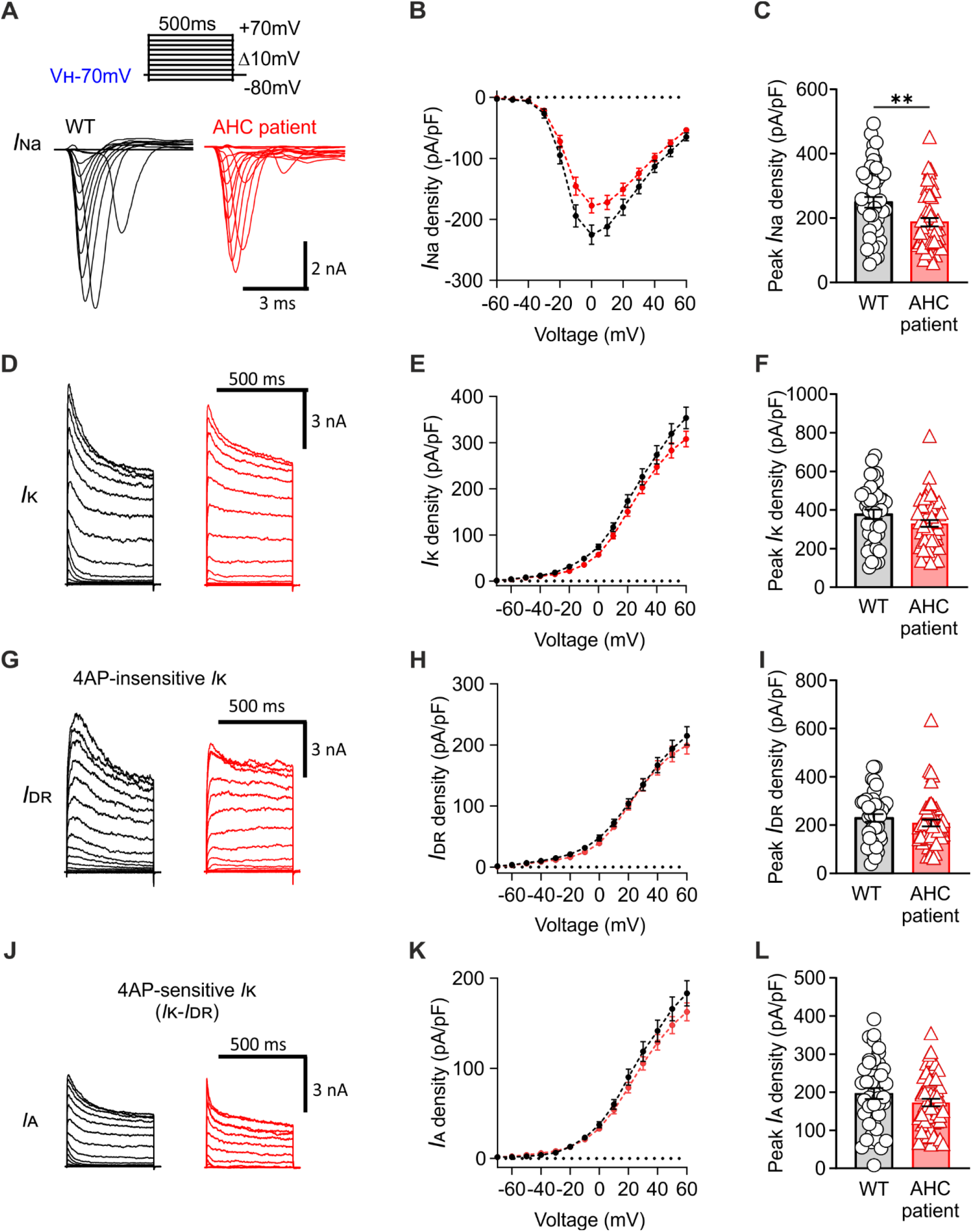
Voltage-dependent sodium currents, but not potassium currents, are reduced in AHC patient-derived iPSC neurons. **(A)** Representative traces of voltage-dependent sodium currents in WT and AHC patient neurons. **(B)** Average current-voltage (I-V) relationship of sodium currents. **(C)** Average peak sodium current density (WT: n = 47; AHC patient: n = 46). **(D)** Representative traces of voltage-dependent potassium currents. **(E)** I-V relationship of total potassium currents. **(F)** Average peak potassium current density (WT: n = 41; AHC patient: n = 51). **(G)** Representative traces of delayed rectifier potassium currents. **(H)** I-V relationship of delayed rectifier potassium currents. **(I)** Average delayed rectifier potassium current density (WT: n = 41; AHC patient: n = 51). **(J)** Representative traces of A-type potassium currents. **(K)** I-V relationship of A-type potassium currents. **(L)** Average A-type potassium current density (WT: n = 41; AHC patient: n = 51). Data are presented as mean ± SEM. *p<0.05.

Potassium currents were separated into delayed-rectifier and fast inactivation (A-type) potassium currents. No significant differences were found in the I-V curve or the maximum potassium current density between patient and control neurons (healthy group: 378.3 ± 23.87 pA/pF, n = 41; patient group: 330.3 ± 18.01 pA/pF, n = 51; P = 0.1050) (Figure 6D-F). Both the delayed-rectifier (DR) potassium current density (healthy group: 229.8 ± 15.76, n = 41; patient group: 209.0 ± 14.51, n = 51; P = 0.3364) (Figure 6G-I) and the A-type potassium current density (healthy group: 196.3 ± 14.78, n = 41; patient group: 172.7 ± 10.18, n = 51; P = 0.1790) (Figure 6J-L) were unaffected by the mutation in the α3 subunit.

Taken together, the reduced AP firing rate, the slowed depolarization phase, and the diminished sodium current density in AHC patient-derived neurons suggest that impaired ion homeostasis, rather than intrinsic alterations in sodium channel dynamics, is the primary contributor to the observed changes in neuronal excitability.

## Discussion

Heterozygous missense mutations in the *ATP1A3* gene, most notably D801N, E815K, and G947R, constitute the most common pathogenic variants associated with AHC, collectively accounting for approximately 60% of all reported cases (Heinzen et al. 2012, Rosewich et al. 2012, Ishii et al. 2013, Viollet et al. 2015). AHC exhibits a wide spectrum of clinical manifestations, with the E815K mutation consistently associated with the most severe neurological, motor, and cognitive impairments. Due to its pronounced phenotypic severity, the E815K variant has become a primary focus of both clinical characterization and mechanistic research.

In this study, iPSC-derived neurons from a patient carrying the E815K mutation were used to investigate the functional consequences of this variant and to elucidate its contribution to the pathogenesis of AHC. The α3 isoform of the Na⁺/K⁺-ATPase plays a crucial role in maintaining neuronal excitability and supporting normal brain development. The importance of this isoform is further underscored by the expanding spectrum of ATP1A3-related neurological disorders, which include AHC, Rapid-onset Dystonia-Parkinsonism (RDP, DYT12) (Brashear et al. 2007, Haq et al. 2019), and CAPOS syndrome (Demos et al. 2014) as the most frequently reported phenotypes. Furthermore, α3 dysfunction has been implicated in the pathogenesis of severe cortical malformations, such as polymicrogyria, which manifest with variable severity (Smith et al. 2021).

Although certain *ATP1A3* mutations, such as R901M, L924P, and Q851R, have been linked to severe malformations of the cerebral cortex (Smith et al. 2021), our data demonstrate that iPSCs carrying the E815K mutation are capable of differentiating comparably to controls into mature inhibitory neurons. This was confirmed by the robust expression of established canonical neuronal markers β3-tubulin and MAP2. Furthermore, functional assessments of synaptogenesis and evoked neuronal activity demonstrated that these processes are largely preserved in E815K-mutant neurons. Previous studies have proposed that aberrant subcellular localization or misfolding of the α3 subunit, associated with specific *ATP1A3* mutations, may contribute to the severity of the resulting phenotype (Arystarkhova et al. 2019).

Subcellular localization of the mutant α3 subunit was examined in patient-derived neurons. Immunocytochemical analysis demonstrated that the E815K-mutant α3 subunit exhibited appropriate somato-dendritic distribution, comparable to that observed in wild-type neurons. This pattern aligns with the conserved trafficking and localization mechanisms previously reported in rodent models (Blom et al. 2011, Liebmann et al. 2018), suggesting an evolutionarily preserved α3 subunit sorting process. These findings indicate that neither impaired neuronal maturation nor missorting of the α3 subunit is a primary pathogenic mechanism in E815K-caused AHC, thereby shifting the focus toward functional deficits as key contributors to the observed phenotype.

Accumulating evidence over the past decade indicates that the residual Na⁺/K⁺-ATPase activity of the mutant α3 subunit exhibits minimal, if any, correlation with clinical disease subtype or the severity of neurological symptoms (Heinzen et al. 2012, Weigand et al. 2014, Arystarkhova et al. 2019, Lazarov et al. 2020). Studies employing heterologous expression systems, including COS-7 cells and cell-free assays with recombinant mutant proteins, have demonstrated that the E815K mutation significantly reduces Na⁺/K⁺-ATPase pump activity (Heinzen et al. 2012, Weigand et al. 2014). Consistent with these findings, patient-derived neurons carrying the E815K mutation exhibit marked ATPase dysfunction, with impairments comparable to those observed in neurons expressing the disease-associated G947R variant (Simmons et al. 2018). Importantly, quantitative analyses revealed that the expression levels of the α3 subunit in E815K-mutant neurons are comparable to, or even exceed, those observed in wild-type controls (Snow et al. 2020). Moreover, the α3 subunits in the AHC patient neurons traffic normally to the soma and dendrites, indicating that the observed functional deficits arise from impaired pump activity rather than reduced abundance or missorting. In contrast, the α1 subunit maintains normal function in patient-derived neurons, suggesting that its interactions with the β subunits necessary for membrane insertion and stabilization are preserved in the mutant neurons.

Electrophysiological analyses revealed that E815K neurons exhibit a depolarized resting membrane potential and reduced excitability, consistent with disrupted ion homeostasis due to the α3 subunit dysfunction. These findings align with previous studies on AHC-associated mutations, including the G947R variant (Simmons et al. 2018). However, in contrast to G947R, where reduced excitability has been linked to decreased availability of voltage-gated sodium channels, our findings indicate that the predominant mechanism in E815K-mutant neurons is a reduced sodium electrochemical driving force, which consequently compromises neuronal excitability.

The α3 subunit represents the predominant Na⁺/K⁺-ATPase isoform in inhibitory neurons, including PV⁺ interneurons and Purkinje cells across multiple brain regions, underscoring its critical role, relative to α1, in the maintenance of resting membrane stability (Richards et al. 2007, Anderson et al. 2010, Murata et al. 2020). Consequently, α3 dysfunction, whether due to G947R or E815K mutations, is likely to impair inhibitory neuron activation, promoting network hyperexcitability and pathological synchronization. This aligns with observed elevated rates of epileptogenesis observed in AHC mouse models (ATP1A3^+/E815K^, ATP1A3^+/D801^) (Hunanyan et al. 2015, Helseth et al. 2018), and the increased seizure susceptibility in patients (Rosewich et al. 2012, Uchitel et al. 2019), providing direct evidence of brain-wide hyperexcitability. Thus, the GABAergic dysfunction, manifesting as an imbalance between excitatory and inhibitory neurotransmission, may underlie the diverse spectrum of neurological and psychiatric symptoms observed in AHC.

In summary, our study elucidates that the *ATP1A3* E815K mutation disrupts neuronal function through a specific biophysical mechanism associated with an impaired sodium electrochemical gradient. This selectively destabilizes inhibitory circuits, leading to network hyperexcitability. While the mutation does not impair neuronal maturation and α3 subunit trafficking, it leads to depolarized resting membrane potentials and reduced excitability, contribution to the excitatory-inhibitory imbalances that is also observed in other AHC-associated variants such as G947R. These findings not only deepen our understanding of AHC pathophysiology but also provide a mechanistic link between α3 Na^+^/K^+^ ATPase dysfunction and a broader spectrum of related neurological disorders, including epilepsy and dystonia. By highlighting the particular vulnerability of inhibitory neurons to the E815K mutation, our work underscores the need for targeted therapeutic strategies aimed at restoring ionic homeostasis. Future investigations using patient-derived brain organoids will enable a direct assessment of the balance between excitatory and inhibitory neurons as well as network oscillatory activity. Furthermore, studies on astrocyte-mediated compensation and mutation-specific pharmacotherapies will be essential to translate these mechanistic insights into clinical treatment approaches for this devastating early-onset neurodevelopmental disorder.

## Supporting information

Supplementary Full western blots

## Acknowledgements

We acknowledge the great contributions of Dr. Elenor Lazarov to this project, who sadly passed away before the completion of the manuscript. We thank Corinna Dickel for expert technical assistance.

## Funding statement

The study was supported by the German Research Foundation (DFG; RO 6308/2-1) and the German Patient Advocacy Group AHC Deutschland e.V. (https://ahc-kids.de/) to H.R.; by the DFG Collaborative Research Center TRR274 and the DFG under Germany’s Excellence Strategy (EXC 2067/1-390729940) to J.G.; and by the German Center for Child and Adolescent Health (DZKJ; Federal Ministry of Research, Technology and Space [BMFTR], 01GL2402) to J.G., MP.Z., and H.R.

## Competing interests

The authors declare that they have no competing interests.

## Author contributions

**M.M.** Data curation, writing - original draft and visualization, writing - review and editing; **C.L.** Investigation, data curation and formal analysis; **E.L.** Investigation, data curation and formal analysis; **HJ.R.** Methodology; **A.O.** Investigation and data curation; **L.C.** Methodology, data curation and formal analysis; **M.P.Z.** Methodology; **J.G.** Conceptualization, Funding Acquisition, Writing - review & editing; **JS.R.** Writing - review & editing, Supervision, Methodology; **H.R.** Conceptualization, Funding Acquisition, Methodology, Project Administration, Resources, Supervision, Validation, Writing - review & editing.

## Data availability

The data that support the findings of this study are available from the corresponding author, upon reasonable request.

## Declaration of generative AI and AI-assisted technologies

During the preparation of this work, the authors used AI-assisted tools only for language editing, text organization, and clarity improvements. After using these tools, the authors carefully reviewed and edited all content and take full responsibility for the scientific accuracy and integrity of the manuscript.

---

**Figure S1.**
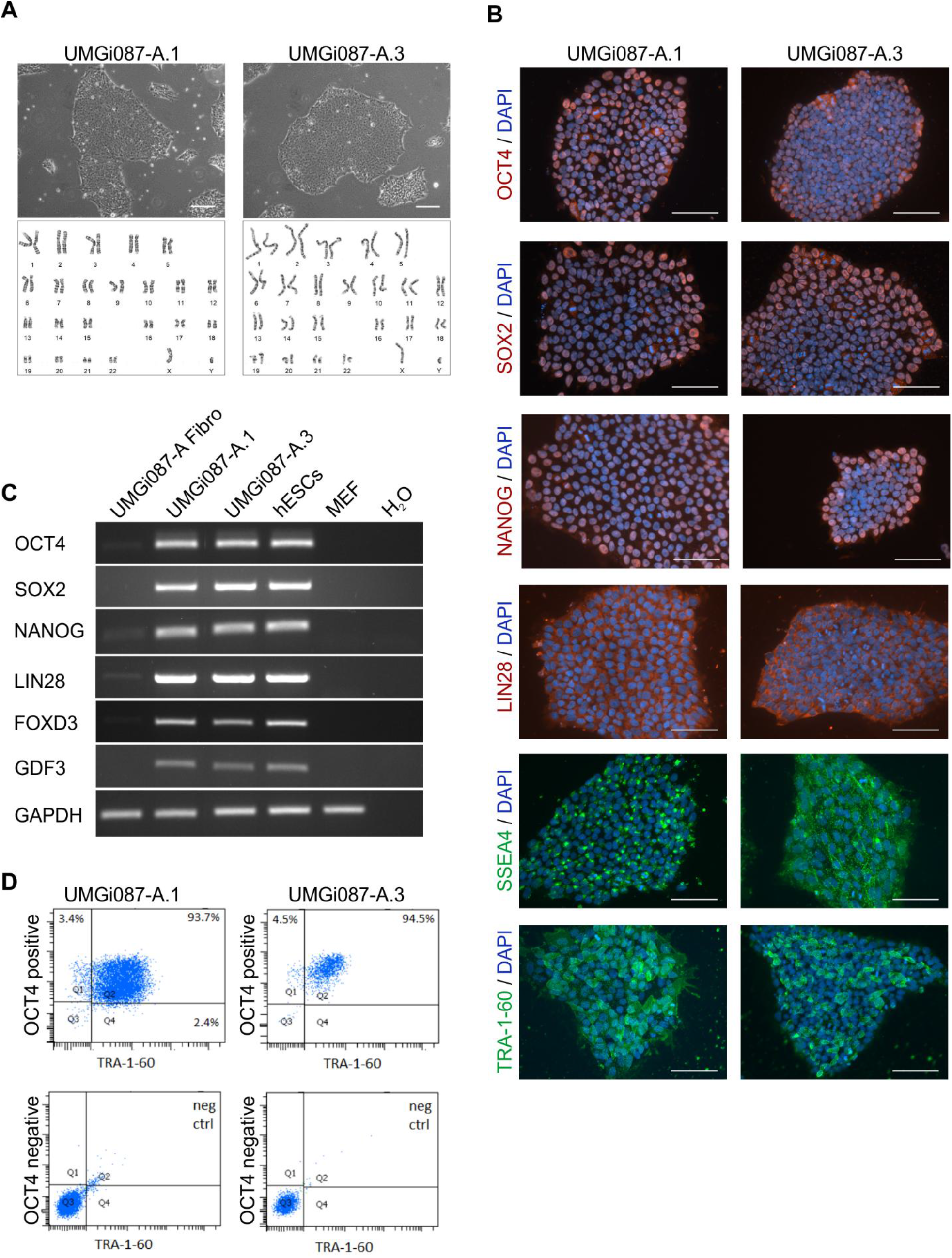
Characterization of the AHC patient derived original iPSC lines (UMGi087-A.1 and -A.3) used in this study. **(A)** Upper panel: Representative images of iPSC colonies from both lines in phase-contrast, showing typical morphology of human pluripotent stem cells (scale bar 200µm). Lower panel: Both iPSC lines exhibit a normal karyotype as assessed by G-band analysis. **(B)** iPSC clones show expression of typical pluripotency markers (OCT4, SOX2, NANOG, LIN28, SSEA4, TRA-1-60) by immunocytochemistry staining (scale bar 100µm). **(C)** The expression of pluripotency markers (OCT4, SOX2, NANOG, LIN28, FOXD3, GDF3) in iPSC lines compared to primary cultures of dermal fibroblasts (UMGGi087-A Fibro) was confirmed by reverse transcriptase-PCR. Human embryonic stem cells (hESCs) and mouse embryonic fibroblasts (MEF) were used as positive and negative controls, respectively. **(D)** Representative FACS analysis of the markers Oct4 and TRA-1-60 was done during line maintenance to validate the pluripotency in both AHC patient derived lines.

## Notes

### Competing Interest Statement

The authors have declared no competing interest.

