## Supplementary Full western blots for "ATP1A3 E815K Disrupts Sodium Homeostasis and Excitability in Human Alternating Hemiplegia of Childhood Neurons"

Figure S2

A

Full blot for Figure 2 C; AHC, UMGi087-A.1

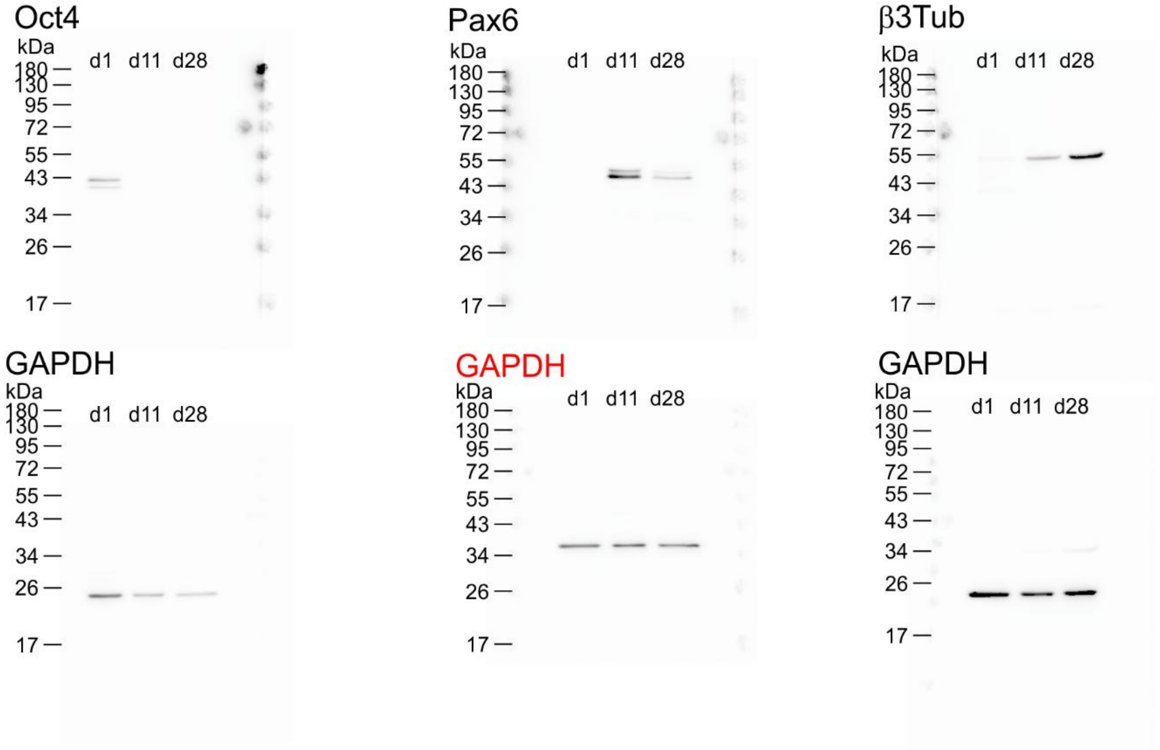


B

Full blot for Figure 2 C; Control, UMGi001-A.1

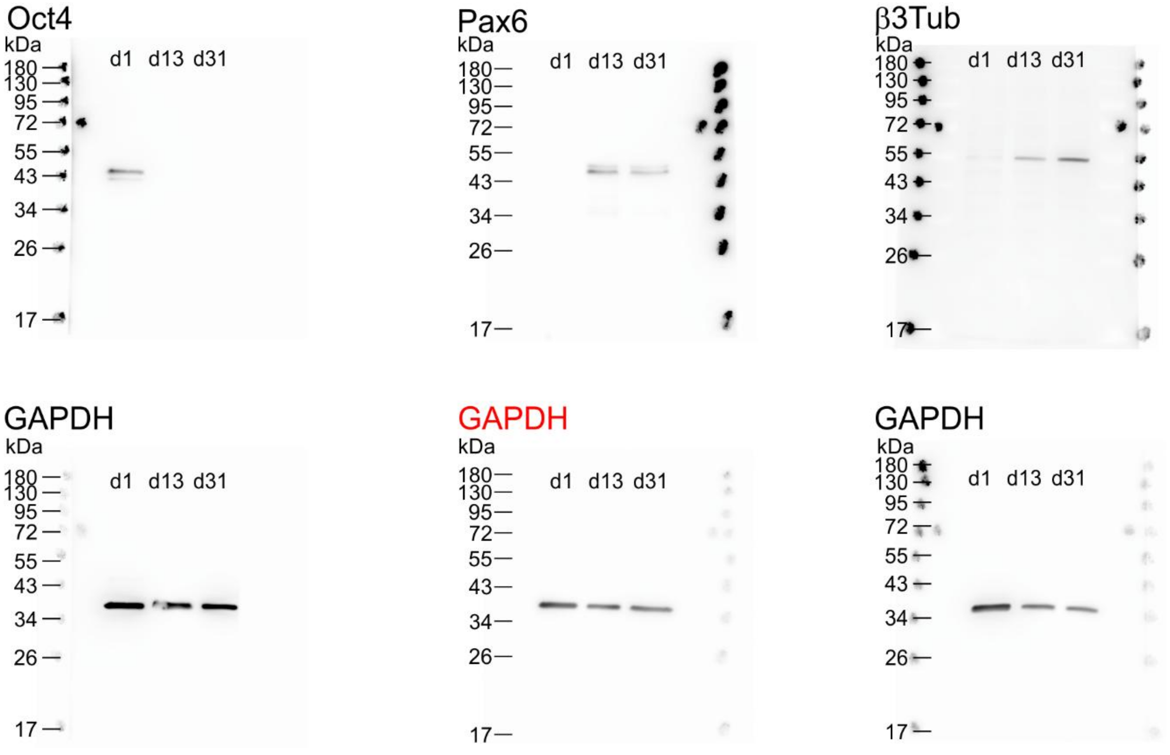

C

Full blot for Figure 2 C; Control, UMGi014-C.14

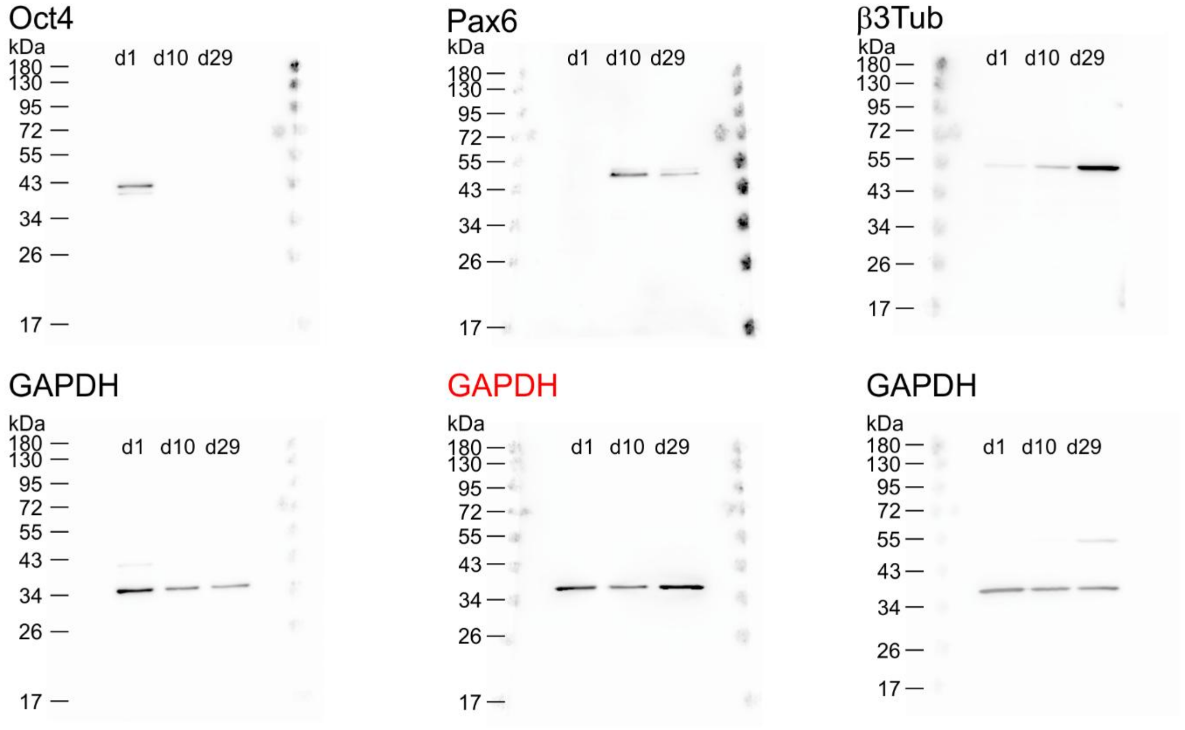


D

Full blot for Figure 2 D

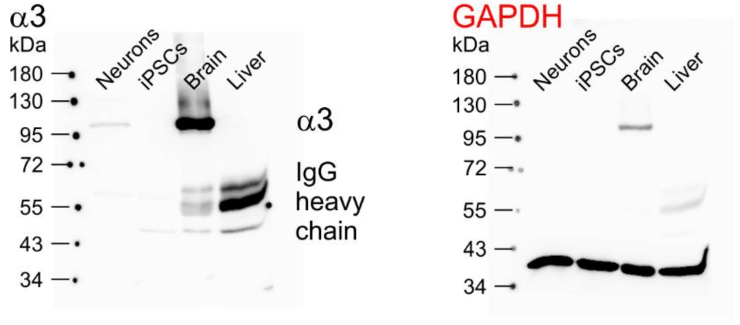


Figure S2: Uncropped and unprocessed blots for Oct4, Pax6, β3-tubulin, and $Na^+/K^+$-ATPase α3 are shown together with the corresponding GAPDH loading controls. Target proteins were detected first, after which membranes were stripped and reprobed for GAPDH. Lane identities and molecular weight markers are indicated on each blot. GAPDH for Pax6 (red) was used as representative GAPDH signals in Figure 2.
