## Supplementary materials tables for "ATP1A3 E815K Disrupts Sodium Homeostasis and Excitability in Human Alternating Hemiplegia of Childhood Neurons"

**Table S1. Antibodies and dilutions used for immunocytochemistry (ICC), flow cytometry (FC)**

**and western blot (WB).**

| Primary antibody | Application | Provider |
|---|---|---|
| ATP1A3, Mouse monoclonal | ICC 1:100<br>WB 1:1,000 | Santa Cruz 374050 |
| βIII-Tubulin TUJ1, Mouse monoclonal | WB 1:2,000<br>ICC 1:250 | MAB1637 |
| DAPI | ICC 1:2000 | Invitrogen D1306 |
| GABA, Rabbit polyclonal | ICC 1:500 | Sigma a2052 |
| GAPDH, Mouse monoclonal | WB 1:10,000 | Abcam 8245 |
| Hoechst 33342 | FC 1:2,000 | Thermo Fisher H3570 |
| LIN28, Goat polyclonal | ICC 1:300 | R&D Systems AF3757 |
| MAP2, Chicken polyclonal | ICC 1:1000 | Novus Biologicals NB300-213 |
| Nanog, Goat polyclonal | ICC 1:200 | R&D Systems AF1997 |
| OCT4, Goat polyclonal | ICC 1:40 | R&D Systems AF1759 |
| OCT3/4, Alexa 647 conjugated, Mouse | FC 1:50<br>WB 1:500 | BD 560329 |
| Pax6, Mouse monoclonal<br>Pax6, Rabbit polyclonal | WB 1:500<br>WB 1:2,000 | Santa Cruz 81649<br>Covance PRB-278P |
| SHANK2, Guinea pig polyclonal | ICC 1:250 | Synaptic Systems 102005 |
| Sox2, Mouse monoclonal | ICC 1:50 | R&D Systems 245610 |
| SSEA4, Mouse monoclonal | ICC 1:100 | Abcam Ab16287 |
| Synapsin1/2, Rabbit polyclonal | ICC 1:500 | Synaptic Systems 106002 |
| TRA-1-60, Mouse monoclonal | ICC 1:200 | Abcam Ab16288 |
| TRA-1-60, Alexa 488 conjugated, mouse | FC 1:50 | BD 560173 |

**Table S2. List of reagents**

| Reagent / Product | Provider | Catalog # |
|---|---|---|
| Accutase | Thermo Fisher | A1110501 |
| AraC | Sigma | C1768 |
| B27 supplement (with vit.A) | Thermo Fisher | 17504044 |

| | | |
|---|---|---|
| Collagenase IV (200 U/ml) | Worthington | LS004188 |
| Dimethyl sulfoxide (DMSO) | Serva | 39757.02 |
| Dispase II | Roche | 04942078001 |
| DMEM low glucose | Thermo Fisher | 11054020 |
| DMEM/F12 Glutamax | Thermo Fisher | 31331028 |
| Dulbecco's Phosphate Buffered Saline (PBS) | Sigma | D8537 |
| Essential 8 medium | Thermo Fisher | A1517001, A1517101 |
| Ethylenediaminetetraacetic acid (EDTA) | AppliChem | A4892,0500 |
| FGF2 | Peprotech | 100-18B |
| Geltrex LDEV free hESC quality | Thermo Fisher | A1413302 |
| Glutamax | Thermo Fisher | 35050061 |
| Insulin | Thermo Fisher | 12585014 |
| Knockout serum replacement | Thermo Fisher | 10828028 |
| LDN 193189 | Selleckchem<br>Axon Medchem | S2618<br>Axon1509 |
| L-glutamine | Thermo Fisher | 25030081 |
| Matrigel | BD | 354230 |
| 2-Mercaptoethanol | Sigma | M3148 |
| Neurobasal medium | Thermo Fisher | 21103049 |
| Non-essential amino acids (NEAA) | Sigma | M7145 |
| N2 supplement | Thermo Fisher | 17502048 |
| Prolong Gold antifade with DAPI | Thermo Fisher | P36935 |
| Rock inhibitor Y-27632 dihydrochloride | Tocris | 1254 |
| SB-431542 | Selleckchem | S1067 |
| Sodium chloride | Roth | 3957.1 |

**Table S3. Primer sequences, annealing temperatures and cycles used for RT-PCR.**

| Gene | Primer (for/rev) | Length (bps) | Temp / Cycles |
|---|---|---|---|
| FOXD3 | GTGAAGCCGCCTTACTCGTAC / CCGAAGCTCTGCATCATGAG | 353 | 61° / 38 |

| | | | |
|---|---|---|---|
| GAPDH | AGAGGCAGGGATGATGTTCT / TCTGCTGATGCCCCCATGTT | 258 | 60° / 30 |
| GDF3 | TTCGCTTTCTCCCAGACCAAGGTTTC / TACATCCAGCAGGTTGAAGTGAACAGCACC | 311 | 54° / 32 |
| LIN28 | AGTAAGCTGCACATGGAAGG / ATTGTGGCTCAATTCTGTGC | 410 | 52° / 36 |
| NANOG | AGTCCCAAAGGCAAACAACCCACTTC / ATCTGCTGGAGGCTGAGGTATTTCTGTCTC | 164 | 64° / 36 |
| OCT4 | GACAACAATGAAAATCTTCAGGAGA / TTCTGGCGCCGGTTACAGAACCA | 218 | 59° / 36 |
| SOX2 | ATGCACCGCTACGACGTGA / CTTTTGCACCCCTCCCATTT | 437 | 60° / 30 |

**Table S4. Primers for genotyping**

| Primer Name | Primer Sequence |
|---|---|
| *ATP1A3* Exon 17-18 PCR Forward Primer | AAAAAAAAAAGGACGTTGGATGA |
| *ATP1A3* Exon 17-18 PCR Reverse Primer | CAGATAGCTCACTGGTTGGTCC |
